# nipalsMCIA: Flexible Multi-Block Dimensionality Reduction in R via Non-linear Iterative Partial Least Squares

**DOI:** 10.1101/2024.06.07.597819

**Authors:** Max Mattessich, Joaquin Reyna, Edel Aron, Ferhat Ay, Misha Kilmer, Steven H. Kleinstein, Anna Konstorum

## Abstract

**Motivation:** With the increased reliance on multi-omics data for bulk and single cell analyses, the availability of robust approaches to perform unsupervised analysis for clustering, visualization, and feature selection is imperative. Joint dimensionality reduction methods can be applied to multi-omics datasets to derive a global sample embedding analogous to single-omic techniques such as Principal Components Analysis (PCA). Multiple co-inertia analysis (MCIA) is a method for joint dimensionality reduction that maximizes the covariance between block- and global-level embeddings. Current implementations for MCIA are not optimized for large datasets such such as those arising from single cell studies, and lack capabilities with respect to embedding new data.

**Results:** We introduce `nipalsMCIA`, an MCIA implementation that solves the objective function using an extension to Non-linear Iterative Partial Least Squares (NIPALS), and shows significant speed-up over earlier implementations that rely on eigendecompositions for single cell multi-omics data. It also removes the dependence on an eigendecomposition for calculating the variance explained, and allows users to perform out-of-sample embedding for new data. `nipalsMCIA` provides users with a variety of pre-processing and parameter options, as well as ease of functionality for down-stream analysis of single-omic and global-embedding factors.

**Availability:** `nipalsMCIA` is available as a BioConductor package at https://bioconductor.org/packages/release/bioc/html/nipalsMCIA.html, and includes detailed documentation and application vignettes. Supplementary Materials are available online.

## 1. Introduction

Multiple co-inertia analysis (MCIA) is a member of the family of joint dimensionality reduction (jDR) methods that extend unsupervised dimension reduction techniques such as Principal Components Analysis (PCA) and Non-negative Matrix Factorization (NMF) to datasets with multiple data *blocks* (alternatively called *views*) [1, 2]. Such methods, also known as multi-block or multi-view analysis algorithms, are becoming increasingly important in the field of bioinformatics, where data is often collected simultaneously using multiple *omics* technologies such as transcriptomics, proteomics, epigenomics, metabolomics, etc. [3].

Here, we present a new implementation in R/Bioconductor of MCIA, `nipalsMCIA`, that uses an extension with proof of monotonic convergence of Non-linear Iterative Partial Least Squares (NIPALS) to solve the MCIA optimization problem [4]. This implementation shows significant speed-up over existing Singular Value Decomposition (SVD)-based approaches for MCIA [5, 6] on large datasets. Furthermore, nipalsMCIA offers users several options for pre-processing and deflation to customize algorithm performance, methodology to perform out-of-sample global embedding, and analysis and visualization capabilities for efficient results interpretation. We show application of `nipalsMCIA` to both bulk and single cell multi-omics data. The overall workflow that includes the optimization steps and analyses for `nipalsMCIA` is outlined in Figure 1.

**Figure 1.**
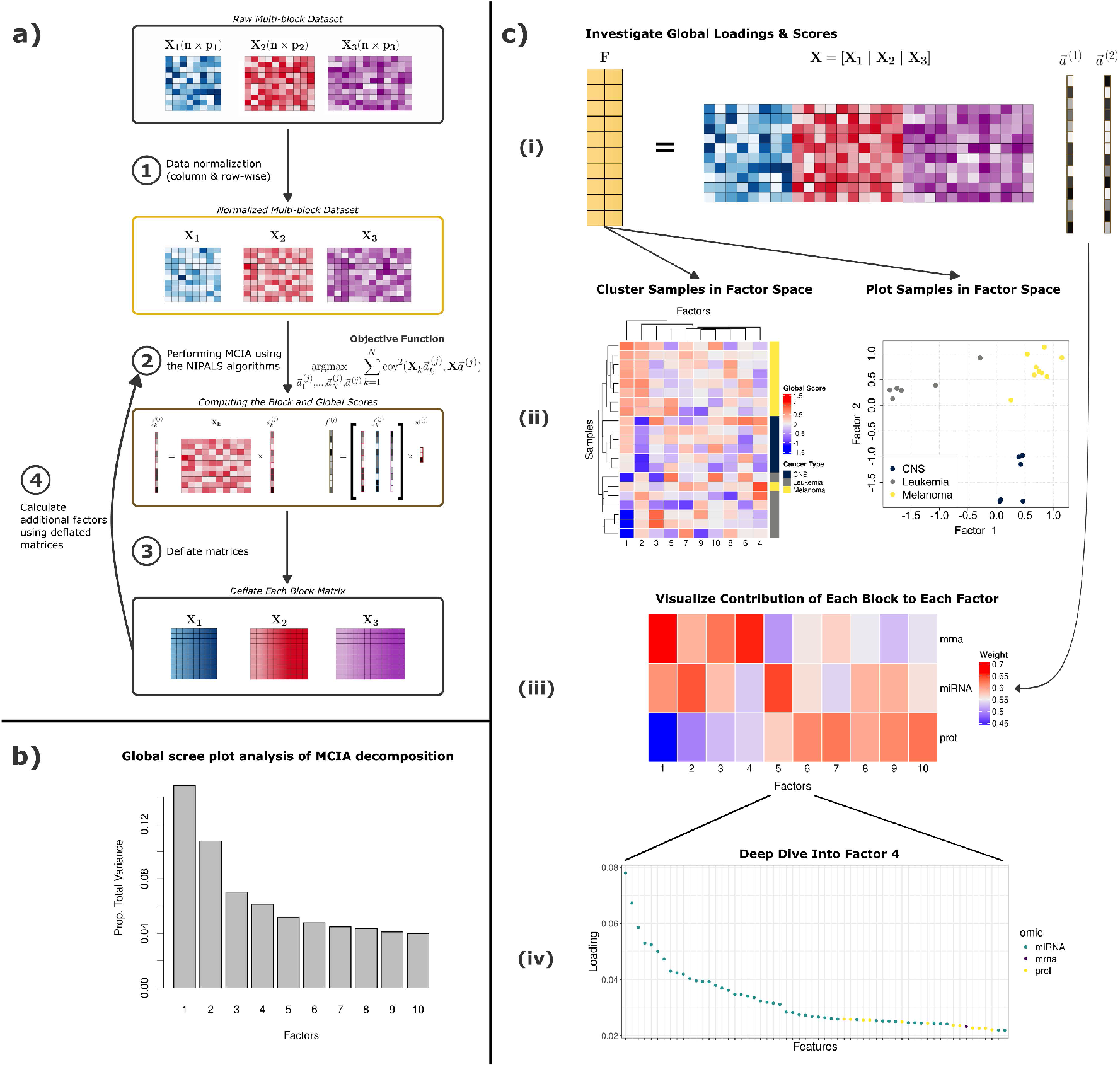
Workflow overview for `nipalsMCIA` performed on the three-block NCI60 data from the main text. a) A breakdown of the NIPALS algorithm for performing MCIA. Data blocks are normalized before scores and loadings are computed to satisfy the objective function. Higher-order results are then computed after the data has been deflated with the current scores or loadings. b) Scree plot for the proportion of variance explained by each order of global score/loading. c) Scheme for interpreting the global loadings and scores. (i) Global scores are calculated from the global data matrix and global loadings. (ii) Global scores represent low-dimensional embeddings of the data used to cluster samples via hierarchical clustering. Colors represent the three different cancer types associated with each sample (iii) Block contributions vectors plotted to visualize the weight of each block on each order of global score. (iv) The first global loadings vector is plotted to identify the top features for the first global score.

## 2. MCIA: theoretical background

### 2.1 Notation and preliminaries

Scalars, vectors, and matrices are represented in lowercase script (a), lowercase script with a vector symbol 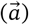 and bold uppercase script (**A**), respectively. The *i*th column vector of a matrix **A** is denoted 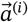.

Since we are evaluating several datasets (termed *blocks*) simultaneously, the sample-by-feature data matrix for the *k*th block is labeled as **X**_*k*_. We denote the column-wise concatenation of data blocks as the ‘global’ data matrix **X** = [**X**_1_|…|**X**_*N*_].

### 2.2 Loadings and Scores

MCIA extends the concept from PCA of deriving principal components (which we term *scores*) and loadings (which we also term *loadings*) to the multi-block setting. The loadings are a set of optimal axes in feature space, while the scores are the projection coefficients of the samples onto these axes. Unlike PCA, MCIA generates two types of scores and loadings, one set for all the data (*global* scores/loadings), and the other for the individual omics (*block* scores/loadings). The number of scores/loadings generated is equal to the dimension of the MCIA embedding of the data, which we will denote as *R*.

Originally, the optimization criteria for MCIA were presented using the concept of *statistical triplets* [7, 5]. The criteria can equivalently be represented as a parameterized member of the Regularized Canonical Correlation Analysis (RGCCA) family of multi-variate dimension reduction methods [2, 8], which is consistent with the optimization criteria that is solved by an extension of the NIPALS algorithm [4]. We review these criteria below.

In the multi-block dataset, each block must share the same samples (rows), but the number of features (columns) *p*_*k*_ in each block *k* can vary. `nipalsMCIA` generates distinct block-level scores 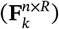 and loadings 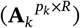, and global scores (**F**^*n*×*R*^) and loadings (**A**^*p*×*R*^), where 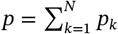, and *R* is the dimension of the embedding. The scores, loadings, and data matrices are related as follows,

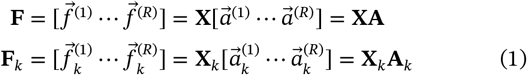

for all blocks *k* = 1, …, *N* (Figure 1c, i).

Scores and loadings are computed by `nipalsMCIA` to satisfy the objective function and orthogonality constraints

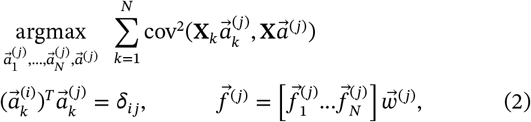

Where 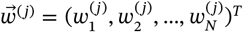 is a vector of block contributions to the j^th^ order glbal score, with constraint 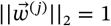 for all orders *j* = 1, …, *R* as in [4], and is the Kronecker delta function. Equation (2) is solved separately for each order (j) up to the dimension of the embedding,*R*. The block scores 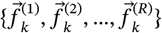 represent a *R*-dimensional embedding of the samples in the orthonormal set of block loadings vectors for block *k*. This contrasts with Consensus PCA (CPCA), which solves for the same objective function as MCIA, but with an orthogonality constraint on the global scores instead of the block loadings [9]. In `nipalsMCIA`, users can choose to use either method.

### 2.3 NIPALS strategy for computing MCIA

Several methods exist for computing MCIA, including direct computation from the principal components of the covariance matrix (see [2]). The implementation in `nipalsMCIA` uses an extension of Nonlinear Iterative Partial Least Squares method (NIPALS) [4]. NIPALS was first introduced as an iterative (power) method to estimate principal components [10, 11], and later extended to the multi-block setting [12]. A modification of the multi-block algorithm was proven to have monotonic convergence [4]. Since the NIPALS procedure is iterative, it does not require a full eigendecomposition. Moreover, it easily enables a choice of deflation methods. In `nipalsMCIA`, the stable multi-block extension to NIPALS [4] is implemented with deflation options for both MCIA and CPCA. Additionally, variance explained by each component is also calculated without reference to an eigendecomposition calculation.

## 3. Usage and functionality

Since MCIA is designed to handle multiple omics data blocks, preprocessing options are available both at within- and whole-block levels. The latter is recommended to account for potential disparities in block size.

### 3.1 Analysis & Interpretation

The `nipals_multiblock` function is used to run MCIA in `nipalsMCIA`. The function outputs an object of the `NipalsResult` class, which includes the global scores and loadings, block scores and loadings, the global score eigenvalues, and the block score contributions vector for all orders up to the maximum specified via the `num_PCs` argument. The global scores represent the projection of the multi-block data in the reduced space, and can be plotted with or without corresponding block scores (Figure 1C, ii). The contribution of each block to the global score can be easily visualized (Figure 1C, iii), along with high-scoring features (Figure 1C, iv).

Vignettes providing full analysis pipelines using `nipalsMCIA` for bulk and single cell data are available with the package. The example bulk data is a subset of the National Cancer Institute 60 tumor-cell line screen (NCI60 data) [13, 8]. It includes RNA-Seq, miRNA, and protein data from 21 cell lines that correspond to three cancer subtypes (brain, leukemia, and melanoma). The single cell data is sourced from 10x Genomics and includes both gene expression and cell surface antibody data [14]. The single cell analysis vignette includes instruction on how to obtain, process, and prepare the dataset for `nipalsMCIA`, along with a demonstration of the capability of `nipalsMCIA` for effectively clustering known cell types in a computationally efficient manner.

### 3.2 Out-of-sample embedding

The loadings vectors generated by MCIA on a dataset **X** represent linear combinations of the original features of **X**. Therefore, after computing MCIA on a training dataset, one can use the associated loadings vectors to predict global embeddings for a test dataset of new observations of the same features. `nipalsMCIA` provides the `predict_gs` function for this task.

This can be valuable for testing the quality of the embedding, as well as embedding new data without rerunning the decomposition. We provide a vignette in the package showing how this can be done using the NCI60 data set, using 70% of the data to train the model, and then deriving global scores for the remaining 30%.

## 4. Computation time comparison for MCIA algorithms

We used three datasets to compare the performance of `nipalsMCIA` with two other implementations of MCIA: MOGSA [6], and Omicade [5]. The three datasets are composed of the NCI60 data, the 10x single cell data filtered for the top 2000 most variable genes, and the same single cell data without filtering. Data pre-processing was standardized across all algorithms and a decomposition for 10 factors was performed across all datasets and implementations. All experiments were performed in R 4.3.0 on a MacBook with 3.2GHz and 16GB RAM. The dimensions of the datasets and performance are shown in Table 1. We observe that while MOGSA has slightly faster performance than `nipalsMCIA` and Omicade on the smaller NCI60 dataset, `nipalsMCIA` is an order of magnitude faster for both the filtered and full single cell datasets, even when using the ‘fast SVD’ option in `MOGSA`. The speedup offered by `nipalsMCIA` thus opens up capabilities for practical deployment of `nipalsMCIA` on a larger variety of datasets, including high-dimensional single cell data.

**Table 1.** Computation time (in seconds) comparison for different MCIA implementations and datasets.

| Implementation | Dataset (dimension) |  |  |
| --- | --- | --- | --- |
|  | NCI60<br>21 x (12895, 547, 7016) | Single cell (filtered)<br>4193 x (2000, 32) | Single cell (full)<sup>1</sup><br>4193 x (33538, 32) |
| `nipalsMCIA` | 2.3 | 15.32 | 289.46 |
| `MOGSA` | 0.53 | 519.55 | NA |
| `MOGSA (fast SVD)` | 0.38 | 434.84 | 13,840.66 |
| `Omicade` | 2.66 | 1089.53 | NA |

## 5. Discussion

The accessibility of next-generation sequencing and other high-throughput biological assays are resulting in an increase of multi-block (or multi-modal) datasets [15, 16, 17, 18]. Analysis of these data are facilitated by the application of joint dimensionality reduction methods such as MCIA. `nipalsMCIA` is a comprehensive R package that implements MCIA in a highly efficient manner using the NIPALS algorithm. The package features various pre-processing and analysis options, is much faster for large input datsets compared with existing packages, supports the projection for out-of-sample scores, and offers visualization options for scores and top-magnitude loadings at each order.

## 6. Supplementary Materials

The Supplementary Materials include additional information on the NIPALS algorithm implemented in `nipalsMCIA`, in-depth discussion of data pre-processing options, detailed overview of the calculations for variance explained and out-of-sample embedding, as well as a summary of the results (detailed more fully in the vignettes) corresponding to out-of-sample embedding of NCI60 data and application of `nipalsMCIA` to single cell data.

## 7. Acknowledgments

This work was supported in part by National Institutes of Health (NIH) grants R21AI176204 to S.H.K. and M.K., R35GM128938 to F.A., and U01AI150753.

## Footnotes

1 Due to the slow performance of `MOGSA` without the fast SVD option and `Omicade`, only `nipalsMCIA` and `MOGSA` with the fast SVD option were tested for the full single cell dataset.

## Notes

### Competing Interest Statement

SHK receives consulting fees from Peraton.

## References

[1] Laura Cantini et al. “Benchmarking joint multi-omics dimensionality reduction approaches for the study of cancer”. en. In: Nature Communications 12.1 (Jan. 2021). Number: 1 Publisher: Nature Publishing Group, p. 124. ISSN: 2041-1723. doi: 10.1038/s41467-020-20430-7. URL: https://www.nature.com/articles/s41467-020-20430-7 (visited on 03/17/2023).

[2] Arthur Tenenhaus and Michel Tenenhaus. “Regularized Generalized Canonical Correlation Analysis”. en. In: Psychometrika 76.2 (Apr. 2011), pp. 257–284. ISSN: 1860-0980. doi: 10.1007/s11336-011-9206-8. URL: https://doi.org/10.1007/s11336-011-9206-8 (visited on 03/17/2023).

[3] Konrad J. Karczewski and Michael P. Snyder. “Integrative omics for health and disease”. eng. In: Nature Reviews. Genetics 19.5 (May 2018), pp. 299–310. ISSN: 1471-0064. doi: 10.1038/nrg.2018.4.

[4] Mohamed Hanafi, Achim Kohler, and El-Mostafa Qannari. “Connections between multiple co-inertia analysis and consensus principal component analysis”. en. In: Chemometrics and Intelligent Laboratory Systems. Multiway and Multiset Data Analysis 106.1 (Mar. 2011), pp. 37–40. ISSN: 0169-7439. doi: 10.1016/j.chemolab.2010.05.010. URL: https://www.sciencedirect.com/science/article/pii/S0169743910000869 (visited on 03/17/2023).

[5] Chen Meng et al. “A multivariate approach to the integration of multi-omics datasets”. In: BMC Bioinformatics 15.1 (May 2014), p. 162. ISSN: 1471-2105. doi: 10.1186/1471-2105-15-162. URL: https://doi.org/10.1186/1471-2105-15-162 (visited on 04/01/2023).

[6] Chen Meng et al. “MOGSA: Integrative Single Sample Geneset Analysis of Multiple Omics Data”. In: Molecular & Cellular Proteomics: MCP 18.8 Suppl 1 (Aug. 2019), S153–S168. ISSN: 1535-9476. doi: 10.1074/mcp.TIR118.001251. URL: https://www.ncbi.nlm.nih.gov/pmc/articles/PMC6692785/ (visited on 04/01/2023).

[7] Daniel Chessel and M Hanafi. “Analyses de la co-inertie de K nuages de points”. In: Revue de statistique appliquée 44.2 (1996), pp. 35–60.

[8] Chen Meng et al. “Dimension reduction techniques for the integrative analysis of multi-omics data”. In: Briefings in Bioinformatics 17.4 (July 2016), pp. 628–641. ISSN: 1467-5463. doi: 10.1093/bib/bbv108. URL: https://doi.org/10.1093/bib/bbv108 (visited on 04/06/2023).

[9] Sahar Hassani et al. “Deflation strategies for multi-block principal component analysis revisited”. In: Chemometrics and Intelligent Laboratory Systems 120 (Jan. 2013), pp. 154–168. ISSN: 0169-7439. doi: 10.1016/j.chemolab.2012.08.011. URL: https://www.sciencedirect.com/science/article/pii/S016974391200189X (visited on 03/13/2024).

[10] Herman Wold. “Nonlinear Estimation by Iterative Least Squares Procedures in: David, FN (Hrsg.), Festschrift for J”. In: Neyman: Research Papers in Statistics, London (1966).

[11] Yoshikatsu Miyashita et al. “Comments on the NIPALS algorithm”. In: Journal of chemometrics 4.1 (1990), pp. 97–100.

[12] Wold, S. et al. “PLS Modeling With Latent Variables in Two or More Dimensions”. In: Frankfurt PLS meeting (1987).

[13] Robert H. Shoemaker. “The NCI60 human tumour cell line anticancer drug screen”. eng. In: Nature Reviews. Cancer 6.10 (Oct. 2006), pp. 813–823. ISSN: 1474-175X. doi: 10.1038/nrc1951.

[14] 10x Genomics. “5k Peripheral blood mononuclear cells (PBMCs) from a healthy donor with cell surface proteins (v3 chemistry)”. en. In: (May 2019). URL: https://support.10xgenomics.com/single-cell-gene-expression/datasets/3.0.2/5k_pbmc_protein_v3.

[15] Yasset Perez-Riverol. “Discovering and Linking Public ‘Omics’ Datasets using the Omics Discovery Index”. eng. In: Nature Biotechnology (May 2017). ISSN: 1546-1696. doi: 10.1038/nbt.3790.

[16] Suhas V. Vasaikar. “LinkedOmics: analyzing multi-omics data within and across 32 cancer types”. eng. In: Nucleic Acids Research (Jan. 2018). doi: 10.1093/nar/gkx1090.

[17] Ana Conesa. “Making multi-omics data accessible to researchers”. eng. In: Scientific data (Oct. 2019). doi: doi: 10.1038/s41597-019-0258-4.

[18] Pedrum Mohammadi-Shemirani. “From ‘Omics to Multiomics Technologies: the Discovery of Novel Causal Mediators”. In: Current atherosclerosis reports (Jan. 2023). doi: doi: 10.1007/s11883-022-01078-8.

