## Supplementary Materials for "nipalsMCIA: Flexible Multi-Block Dimensionality Reduction in R via Non-linear Iterative Partial Least Squares"

#### S1 NIPALS strategy for computing MCIA and CPCA

`nipalsMCIA` computes scores and loadings for two closely-related methods, Multiple Co-Inertia Analysis (MCIA) and Consensus PCA (CPCA). Both satisfy the objective function

$$\underset{\vec{a}_1^{(j)},\ldots,\vec{a}_N^{(j)},\vec{a}^{(j)}}{\operatorname{argmax}} \sum_{k=1}^{N} \operatorname{cov}^2(\mathbf{X}_k^{(j)}\vec{a}_k^{(j)}, \mathbf{X}^{(j)}\vec{a}^{(j)}) \tag{1}$$

for all blocks $k = 1, ..., N$ and where $\mathbf{X}^{(j)} = \left[X_1^{(j)}|...|X_N^{(j)}\right]$ is the data matrix at any deflation step $j = 1, ..., R$. Equation (1) is solved separately for each order $(j)$ of scores/loadings generated, where block loadings of different orders are constrained by orthogonality conditions. For the two options available in `nipalsMCIA`, these constraints are

$$(\vec{a}_k^{(i)})^T\vec{a}_k^{(j)} = \delta_{ij} \quad \text{(MCIA)} \qquad (\vec{f}_k^{(i)})^T\vec{f}_k^{(j)} = \delta_{ij} \quad \text{(CPCA)} \qquad i, j = 1, ..., R \tag{2}$$

for all blocks $k = 1, ..., N$, where $\delta_{ij}$ is the Kronecker delta function, and $R$ is the maximum order of scores/loadings computed. Note that in the original implementations of MCIA, there is an additional constraint such that $\operatorname{var}(\mathbf{X}^{(j)}\vec{a}^{(j)}) = 1$ [3]. For the NIPALS implementation of MCIA, this constraint is not required [5], adding it only imparts a scaling of each global score [9], and does not impact the calculation of the subsequent global scores. Moreover, without the constraint, calculation of variance explained for each order can be associated directly with the variance of the global score (see Section S3).

In `nipalsMCIA`, the implementation of the NIPALS algorithm is based on [4, 5]. It begins with a random starting vector and, through successive projections onto the data and scores vectors, converges to the solution of the optimality criterion (Equation (1)). Algorithm 1 illustrates the iteration process to generate one order of block and global scores/loadings vectors. The iteration process is stopped when the change in the covariance criterion from Equation (1) is below a specified tolerance, i.e.

$$|C_{\text{current}} - C_{\text{prev}}| < \mathbf{tol},$$

where $C \equiv \sum_{k=1}^{N} \operatorname{cov}^2(\vec{f}_k, \vec{f})$ is compared at the current and previous steps of the NIPALS iteration.

The orthogonality condition in (2) is enforced through a *deflation step*, whereby all data aligned with previously-computed block loadings is subtracted from the blocks. Given a set of block loadings

**Algorithm 1** Steps for the iteration stage of the modified NIPALS algorithm. Adapted from [4].

0.) Let $\vec{f} \in \mathbb{R}^{n\times 1}$ be an arbitrary starting vector

1.) Compute block loadings $\vec{a}_k = \frac{\mathbf{X}_k^T \vec{f}}{\vec{f}^T \vec{f}}$ for each block $k = 1, ..., N$

2.) Normalize the block loadings to $\vec{a}_k^T \vec{a}_k = 1$ for $k = 1, ..., N$

3.) Compute block scores $\vec{f}_k = \mathbf{X}_k \vec{a}_k$ for $k = 1, ..., N$

4.) Create an $n \times N$ matrix of block scores $\mathbf{T} = [\vec{f}_1|...|\vec{f}_N]$

5.) Compute global weight vectors $\vec{w} = \frac{\mathbf{T}^T \vec{f}}{\vec{f}^T \vec{f}}$

6.) Normalize the global weight vectors to $\vec{w}^T \vec{w} = 1$ for $k = 1, ..., N$

7.) Set starting vector to $\vec{f} = \mathbf{T}\vec{w}$ and return to step 1.)

Repeat until the global and block scores converge in the chosen convergence metric.

of rank $j$, $\{\vec{a}_1^{(j)}, ..., \vec{a}_N^{(j)}\}$, the deflated blocks are computed via

$$\mathbf{X}_k^{(j+1)} = \mathbf{X}_k^{(j)} - \mathbf{X}_k^{(j)} \vec{a}_k^{(j)} \left(\vec{a}_k^{(j)}\right)^T \quad \text{(MCIA)} \qquad \mathbf{X}_k^{(j+1)} = \mathbf{X}_k^{(j)} - \vec{f}^{(j)} \left(\vec{f}^{(j)}\right)^T \mathbf{X}_k^{(j)} \quad \text{(CPCA)}$$

for all blocks $k = 1, .., N$. Scores and loadings of the $(j+1)^{th}$ order can then be computed by replacing the data blocks $\mathbf{X}_k^{(j)}$ with their deflated equivalents $\mathbf{X}_k^{(j+1)}$ in the iteration loop of Algorithm 1.

### S2 Data pre-processing

Within-block pre-processing options include feature-level mean-centering and, optionally, normalization to unit-variance. The default within-block pre-processing method is the *column-profile* option, which offsets the data so it is non-negative, then transforms each entry $x_{ij}$ of the data block $\mathbf{X}_k$ by

$$x_{ij} \mapsto \frac{x_{ij}}{CS_j} - \frac{RS_i}{||\mathbf{X}_k||_1},$$

where $CS_j = \sum_i [\mathbf{X}_k]_{i,j}$ and $RS_i = \sum_j [\mathbf{X}_k]_{i,j}$ represent the sum of the $j^{th}$ column and $i^{th}$ row of $\mathbf{X}_k$ respectively. This reproduces the pre-processing in the `omicade4` package and represents the relative contribution of a variable with respect to the full (block) dataset [6]. This method also results in all features in the dataset having zero mean. These options can be selected when running the `nipals_multiblock` function via the parameter `col_preproc_method`. Note that there is a requirement in MCIA and CPCA that all features must be centered (like with PCA). All available within-block pre-processing options satisfy this requirement.

Whole-block, also termed block-level, pre-processing options include normalization of each block to unit variance, by total number of features, or by the largest singular value (the latter ensures equal contribution of each block to the first global score) [2]. There are no assumptions for block-level pre-processing in MCIA (and there is an option to perform no pre-processing at the block level), but if there is a large disparity in block size, and pre-processing is not performed, larger blocks may dominate algorithm performance. Thus, we recommend taking advantage of the pre-processing options at the block level using the `block_preproc_method` parameter in `nipals_multiblock` (normalization to unit variance is set as default).

### S3 Global Score Variance Explained

#### S3.1 Total Variance

Consider a data matrix $\mathbf{X}_{n\times m}$ of $n$ samples and $m$ variables. We can write $\mathbf{X} = [\vec{x}^1, \vec{x}^2, ..., \vec{x}^m]$. We let $\bar{x}^i = \frac{1}{n}\sum_{j=1}^{m} \vec{x}^i_j$ as the mean of $\vec{x}^i$, $\bar{\mathbf{X}} = [\bar{x}^1, \bar{x}^2, ..., \bar{x}^m]$ the matrix of column means of $\mathbf{X}$ (each column is a constant), and $\mathbf{X}' = \mathbf{X} - \bar{\mathbf{X}}$ the column-centered matrix corresponding to $\mathbf{X}$. For each variable $\vec{x}^i \in \mathbf{X}$, we take $\bar{x}^i$ as the mean of $\vec{x}^i$. Then, the variance of $\vec{x}^i$ is equal to

$$\text{var}(\vec{x}^i) = \frac{1}{n-1}\sum_{j=1}^{n}(\vec{x}^i_j - \bar{x}^i)^2 = \frac{1}{n-1}\vec{x'}^i \cdot \vec{x'}^i, \tag{3}$$

where $\vec{x'}^i$ correspond to the columns of $\mathbf{X}'$.

We define the Total Variance (TV) of a sample-by-variable $(n \times m)$ data matrix $\mathbf{X}$ as follows,

**Definition 1.**

*$TV(\mathbf{X}) = \sum_{i=1}^{m} c_{ii}$, where $c_{ii}$ are the diagonal elements of the covariance matrix of $\mathbf{X}$, $\mathbf{C}_X = \frac{1}{n-1}\mathbf{X}'^T\mathbf{X}'$.*

We note that we can write TV as as the trace of the covariance matrix, $TV = tr(\mathbf{C_X})$. Indeed we use several properties of the trace of a matrix in our discussion, so we will review them here:

**Definition 2.** *The trace of a symmetric $m \times m$ matrix $\mathbf{A}$ is $tr(\mathbf{A}) = \sum_{i=1}^{m} a_{ii}$.*

Two properties of the trace, which we will not prove here, are as follows. For real-valued $m \times m$ matrices $A$, $B$,

$$tr(AB) = tr(BA). \tag{4}$$

And when $B = A^T$,

$$tr(AA^T) = tr(A^TA) = \sum_{j=1}^{m}\sum_{i=1}^{n} a_{ij}^2. \tag{5}$$

We now show that the following definitions for TV are equivalent. These results should be familiar, but they are collected together in a way that can illuminate the relationships between the concepts of TV, PCA, and SVD in order to facilitate the correct application of TV.

**Property 1.** *The following definitions for TV of an $n \times m$ sample-by-variable matrix $\mathbf{X}$ are equivalent,*

*i* $TV(\mathbf{X}) = \sum_{i=1}^{m} c_{ii}$, *where $c_{ii}$ are the diagonal elements of the covariance matrix of $\mathbf{X}$, $\mathbf{C}_X = \frac{1}{n-1}\mathbf{X}'^T\mathbf{X}'$.*

*ii* $TV(\mathbf{X}) = \sum_{i=1}^{m} \lambda_i$ *where $\lambda_i$ are eigenvalues of $\mathbf{C}_X$.*

*iii* $TV(\mathbf{X}) = \sum_{i=1}^{m} s_i^2$, *where $s_i$ are the singular values of the Singular Value Decomposition (SVD) of $\mathbf{X}^* = \frac{1}{\sqrt{n-1}}(\mathbf{X}')$.*

*iv* $TV(\mathbf{X}) = ||\mathbf{X}^*||_F^2$, *where $||\cdot||_F$ is the Frobenius norm.*

*Proof.* We first prove the equivalence of (i) and (ii). Recall that a real symmetric $n \times n$ matrix $\mathbf{A}$ has an eigendecomposition of $\mathbf{A} = \mathbf{PDP}^T$,where $\mathbf{P}$ is orthogonal, and $\mathbf{D}$ is a diagonal matrix whose entries $\{\lambda_i\}_{i=1}^m$ are the eigenvalues of $\mathbf{A}$. By Equation 4,

$$tr(\mathbf{A}) = tr(\mathbf{PDP}^T) = tr(\mathbf{DP}^T\mathbf{P}) = tr(\mathbf{D}). \tag{6}$$

We now observe that if have $\mathbf{A} = \mathbf{C}_X$, then

$$TV(\mathbf{X}) = tr(\mathbf{C}_X) = tr(\mathbf{D}) = \sum_{i=1}^{m} \lambda_i, \tag{7}$$

where $\mathbf{D}$ is the diagonal matrix of the eigendecomposition of $\mathbf{C}_X$.

We now prove the equivalence of (ii) and (iii). The Singular Value Decomposition (SVD) of a matrix $\mathbf{A}$ is the factorization of $\mathbf{A}$, $\mathbf{A} = \mathbf{USV}^T$, where $\mathbf{U}$ and $\mathbf{V}$ are orthonormal, and $\mathbf{S}$ is a diagonal matrix with positive entries. Observing that $(\mathbf{X}^*)^T\mathbf{X}^* = \frac{1}{n-1}(\mathbf{X} - \bar{\mathbf{X}})^T(\mathbf{X} - \bar{\mathbf{X}}) = \mathbf{C}_X$, we consider the SVD of $\mathbf{X}^*$,

$$\mathbf{C}_X = (\mathbf{X}^*)^T\mathbf{X}^* = \mathbf{VS}^T\mathbf{U}^T\mathbf{USV}^T = \mathbf{V}(\mathbf{S}^T\mathbf{S})\mathbf{V}^T. \tag{8}$$

By Equation 8, $(\mathbf{X}^*)^T\mathbf{X}^*$ and $\mathbf{S}^T\mathbf{S}$ are similar, and therefore will have the same eigenvalues. We have shown that that the eigenvalues of $\mathbf{C}_X = (\mathbf{X}^*)^T\mathbf{X}^*$ are the diagonal elements of $\mathbf{D} = \{\lambda_1, \lambda_2, ...\lambda_m\}$. For singular values in the diagonal of $\mathbf{S}$, $\mathbf{S} = \{\sigma_1, \sigma_2, ..., \sigma_m\}$, $\sigma_i^2 = \lambda_i$. By Equation 7,

$$TV(\mathbf{X}) = \sum_{i=1}^{m} \lambda_i = \sum_{i=1}^{m} \sigma_i^2 \tag{9}$$

Finally, we show the equivalence of (i) and (iv). By the property of the trace in Equation 5,

$$||\mathbf{X}^*||_F^2 = \sum_{i,j} \mathbf{X}^{*2}_{i,j} = tr((\mathbf{X})^{*T}\mathbf{X}^*) = TV(\mathbf{X}), \tag{10}$$

since $\mathbf{C}_X = (\mathbf{X}^*)^T(\mathbf{X}^*)$.

□

#### S3.2 Significance of Global Scores

Earlier implementations compute MCIA directly through the singular value decomposition of the global data matrix $\mathbf{X}^{(i)}$ at each deflation step $i$. This follows from the calculation of the $i^{th}$ order global loading $\vec{a}^{(i)}$ as the first principal component loading of the $i^{th}$ order deflated data matrix $\mathbf{X}^{(i)}$ (see [6, 7]). The corresponding eigenvalue of the first principle component is often used as a measure of the significance of that order of global score. Since `nipalsMCIA` does not use an eigendecomposition, we instead link this value directly to the variance of the global score.

**Property 2.**

$$var(\vec{f}^{(i)}) = \lambda^{(i)}, \tag{11}$$

*where* $\vec{f}^{(i)}$ *is the first principal component of* $\mathbf{X}^{(i)}$ *and* $\lambda^{(i)}$ *is the largest eigenvalue of* $\mathbf{C}_X^{(i)}$

*Proof.* We consider the SVD of $\mathbf{X}^{(i)^*}$, $\mathbf{X}^{(i)^*} = \mathbf{U}^i\mathbf{S}^i\mathbf{A}^{i^T}$, where $\mathbf{X}^{(i)^*} = \frac{1}{\sqrt{n-1}}\mathbf{X}^{(i)}$. Since $A^i$ is orthogonal, we can set

$$\mathbf{X}^{(i)^*}\mathbf{A}^i = \mathbf{U}^i S^i \Rightarrow \mathbf{X}^{(i)^*}\vec{a}_1^{(i)} = \sigma_1^{(i)}\vec{u}_1^{(i)} \tag{12}$$

Since $\vec{f}^{(i)} = \mathbf{X}^{(i)}\vec{a}_1^{(i)}$ (note that $\mathbf{X}^{(i)}$ are mean-centered for all $i$, and the columns of $\mathbf{A}^{(i)}$ are the principal component loadings of $\mathbf{X}^{(i)}$, e.g. see [8]), by Property 1,

$$var(\vec{f}^{(i)}) = ||\vec{f}^{(i)^*}||^2 = ||\mathbf{X}^{(i)}\vec{a}_1^{(i)}||^2 = ||\sigma_1^i\vec{u}_1^i||^2 = \sigma_1^{(i)^2} = \lambda^{(i)}, \tag{13}$$

where $\vec{f}^{(i)^*} = \frac{1}{\sqrt{n-1}}(\mathbf{X}^{(i)}\vec{a}_1^{(i)}) = \mathbf{X}^{(i)^*}\vec{a}_1^{(i)}$.

□

Thus, the eigenvalue $\lambda^{(i)}$ is exactly the variance of the global score $\vec{f}^{(i)}$, which is in turn the variance of the data along the global loading vector $\vec{a}^{(i)}$. It is therefore used (e.g. in [6]) as a significance metric for the global scores at each order, similarly to an eigenvalue scree plot in PCA. Since `nipalsMCIA` does not use the singular value decomposition to compute MCIA, we instead directly compute the variance of each global score vector via Equation (3) and report this as the 'eigenvalue' associated with each order of the score. `nipals_multiblock` can also optionally return a scree plot of eigenvalues, which we report as the proportion of variance explained by each order ($PV_i$) by dividing by the total variance of the data,

$$PV_i = \frac{\text{var}\,\vec{f}^{(i)}}{||\mathbf{X}^*||_F^2}, \tag{14}$$

where $||\mathbf{X}^*||_F^2 = TV(\mathbf{X})$ by Property 1.

### S4 Out-of-Sample Embedding with MCIA

It has previously been shown that the embeddings computed via MCIA can be expressed in terms of both the deflated data as well as the original data (e.g. Eq. 12 in [5]). This can be stated by the following property:

**Property 3.** *Any set of order-i scores/loadings vectors* $\left\{\vec{f}^{(i)}, \vec{f}_k^{(i)}, \vec{a}^{(i)}, \vec{a}_k^{(i)} | k = 1, ..., N\right\}$ *computed to satisfy the MCIA criteria satisfy the relations*

$$\vec{f}^{(i)} = \mathbf{X}^{(i)}\vec{a}^{(i)} = \mathbf{X}\vec{a}^{(i)} \qquad \vec{f}_k^{(i)} = \mathbf{X}_k^{(i)}\vec{a}_k^{(i)} = \mathbf{X}_k\vec{a}_k^{(i)} \qquad k = 1, .., N,$$

*where* $\mathbf{X}^{(i)}$ *is the data at the* $i^{th}$ *deflation step and* $\mathbf{X}$ *is the original data.*

*Proof.* Note that by step 7 in Algorithm 1, the $i^{th}$ order global score can be expressed as

$$\vec{f}^{(i)} = \sum_{k=1}^{N} w_k^{(i)}\vec{f}_k^{(i)} = \sum_{k=1}^{N} w_k^{(i)}\mathbf{X}_k^{(i)}\vec{a}_k^{(i)} \tag{15}$$

where $\vec{w}^{(i)} = (w_1^{(i)}, ..., w_N^{(i)})$ is the vector containing the $i^{th}$ order block score weights. Combining (15) with the relation $\vec{f}^{(i)} = \mathbf{X}^{(i)}\vec{a}^{(i)}$ from Equation (1) of the main text, we can similarly express the global loadings $\vec{a}^{(i)}$ as a weighted stacking of the block loadings,

$$\vec{a}^{(i),T} = \left(w_1^{(i)}\vec{a}_1^{(i),T}, \quad w_2^{(i)}\vec{a}_2^{(i),T}, \quad \dots \quad w_N^{(i)}\vec{a}_N^{(i),T}\right). \tag{16}$$

We may now prove the property by induction comparing any sequential orders $0 \leq (i-1) < i$. Given the MCIA deflation method in Section S1,

$$\begin{aligned}\mathbf{X}^{(i)} &= \left(\mathbf{X}_1^{(i)} \;\middle|\; ... \;\middle|\; \mathbf{X}_N^{(i)}\right) \\ &= \left(\mathbf{X}_1^{(i-1)} - \mathbf{X}_1^{(i-1)}\vec{a}_1^{(i-1)}\vec{a}_1^{(i-1),T} \;\middle|\; ... \;\middle|\; \mathbf{X}_N^{(i-1)} - \mathbf{X}_N^{(i-1)}\vec{a}_N^{(i-1)}\vec{a}_N^{(i-1),T}\right).\end{aligned}$$

Applying Equation (16), we obtain

$$\begin{aligned}\left(\mathbf{X}^{(i-1)} - \mathbf{X}^{(i)}\right)\vec{a}^{(i)} &= \left(\mathbf{X}_1^{(i-1)}\vec{a}_1^{(i-1)}\vec{a}_1^{(i-1),T} \;\middle|\; ... \;\middle|\; \mathbf{X}_N^{(i-1)}\vec{a}_N^{(i-1)}\vec{a}_N^{(i-1),T}\right)\vec{a}^{(i)} \\ &= \sum_{k=1}^{N} \mathbf{X}_k^{(i-1)}\vec{a}_k^{(i-1)}\vec{a}_k^{(i-1),T}\left(w_k^{(i)}\vec{a}_k^{(i)}\right) \qquad \text{(by matrix-vector multiplication)} \\ &= \sum_{k=1}^{N} \mathbf{X}_k^{(i-1)}\vec{a}_k^{(i-1)}\left(w_k^{(i)}\vec{a}_k^{(i-1),T}\vec{a}_k^{(i)}\right) = 0\end{aligned}$$

by the orthogonality condition $\vec{a}_k^{(j),T}\vec{a}_k^{(i)} = \delta_{ij} \;\; \forall k = 1, ..., N$. Thus, the global scores can be represented as embeddings in the original data, i.e.

$$\vec{f}^{(i)} = \mathbf{X}^{(i)}\vec{a}^{(i)} = \mathbf{X}^{(i-1)}\vec{a}^{(i)} = ... = \mathbf{X}\vec{a}^{(i)}$$

for any order $i$. An almost identical proof shows that the block scores are also the embeddings of the original data, i.e.

$$\vec{f}_k^{(i)} = \mathbf{X}_k\vec{a}_k^{(i)}, \qquad k = 1, ..., N.$$

□

#### S4.1 Algorithm for out-of-sample embedding using MCIA

`nipalsMCIA` includes functionality to use previously-computed MCIA loadings to predict global scores for new observations of the same $m$ features via the `predict_gs` function. The function requires a `NipalsResult` object computed from MCIA on a pre-processed training dataset $\mathbf{X} = [\mathbf{X}_1|...|\mathbf{X}_N] \in \mathbb{R}^{n\times m}$, containing $R$ orders of block loadings expressed as columns of the matrices $\{\mathbf{A}_k | k = 1, ..., N\}$ along with the corresponding block weights vectors $(\vec{w}^{(1)}, ..., \vec{w}^{(R)})$. If we let $\mathbf{W}_k$ denote the $R \times R$ diagonal matrix with entries $(w_k^{(1)}, ..., w_k^{(R)})$, we can use Property 3 to adapt Equation (15) to apply to a matrix-wise computation of global scores $\mathbf{F}$ from the block scores matrices,

$$\mathbf{F} = \sum_{k=1}^{N} \mathbf{F}_k\mathbf{W}_k = \sum_{k=1}^{N} \mathbf{X}_k\mathbf{A}_k\mathbf{W}_k, \tag{17}$$

where $\mathbf{F}$ and $\mathbf{F}_k$ are the matrices of global scores, block scores, and global loadings as defined in Equation (1) of the main text. Thus given a test dataset $\mathbf{Y} = [\mathbf{Y}_1|...|\mathbf{Y}_N]$ of different samples observed on the same features, we can use Equation (17) and the previously-computed block loadings $\{\mathbf{A}_1, ..., \mathbf{A}_N\}$ to form a matrix $\mathbf{G}$ of predicted global embeddings via Algorithm 2.

#### S4.2 Out-of-sample embedding on the NCI60 data

The NCI60 data was split randomly into a training/test set (70% train/30% test), and `nipals_multiblock` was applied to the training data set. Next, the `predict_gs` function was used to predict the global

**Algorithm 2** Steps for out-of-sample embedding prediction used in `predict_gs()`.

0.) Given inputs from MCIA computed on dataset $\mathbf{X} = [\mathbf{X}_1|...|\mathbf{X}_N]$:

- Prior/training block loadings matrices $\{\mathbf{A}_1, ..., \mathbf{A}_N\}$.
- Prior/training block score weights matrices $\{\mathbf{W}_1, ..., \mathbf{W}_N\}$.
- New/test multi-block data matrix $\mathbf{Y} = [\mathbf{Y}_1|...|\mathbf{Y}_N]$ observed on the same features.

1.) Perform pre-processing on $\mathbf{Y}$ to match all pre-processing used to compute MCIA on $\mathbf{X}$.

2.) Compute the predicted global embeddings matrix for the samples in $\mathbf{Y}$ via

$$\mathbf{G} = \sum_{k=1}^{N} \mathbf{Y}_k \mathbf{A}_k \mathbf{W}_k.$$

scores for the test dataset. Plotting both the training and predicted global for the first two factors (Figure S1), we observe that samples from cell lines of the same cancer types cluster together for both the training and test data. The *Predicting New Scores* vignette provides additional details for this example.

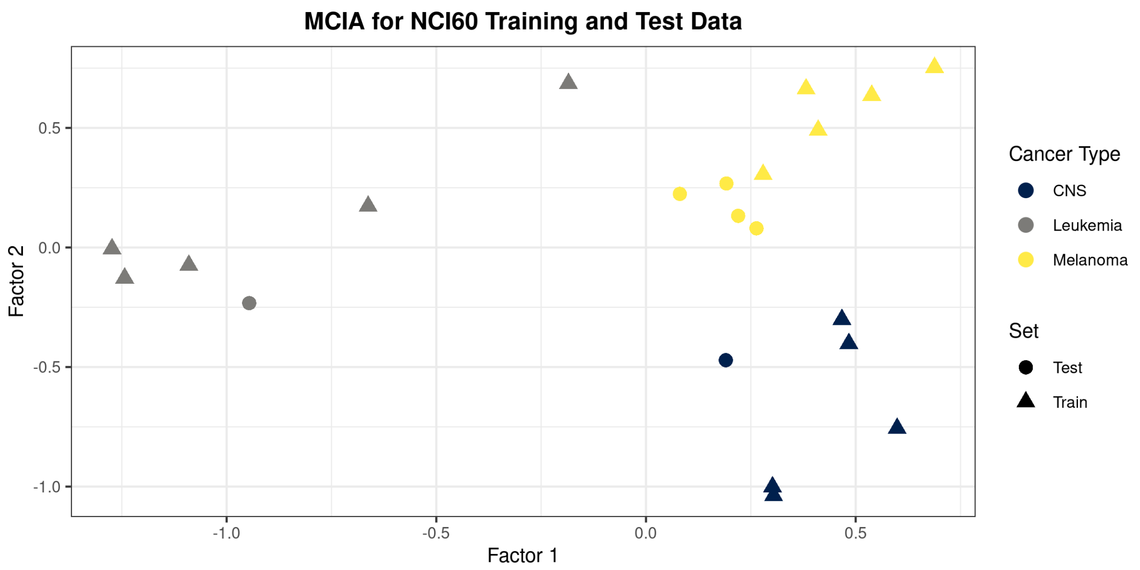


Figure S1: Plotting the global scores for the first two factors of the NCI60 data for both the training (triangle) and test (circle) sets shows that cell types cluster together for both the training and test data.

#### S5 Application of MCIA to a single cell dataset

We demonstrate the utility of `nipalsMCIA` for single cell data in the *Single Cell Analysis* vignette. The example dataset is sourced from 10x Genomics and consists of 5,247 peripheral blood mononuclear cells (PBMCs) from a single healthy donor measured for both gene expression and cell surface antibody abundance ([1]). After performing quality control, the data had dimensions of 4193 (cells) x ((2000 (gene expression) + 32 (antibodies)). Note that original number of 33,538 genes was filtered for the most highly variable genes. We provide details on how to obtain, process, and prepare the dataset as part of the vignette. The vignette begins from the output of the 10x Genomics Cell Ranger pipeline [10], and leads users through steps on how to perform basic quality control (such as the aforementioned filtration for highly variable genes), and cell type annotation prior to generating the input objects for `nipalsMCIA`. Cells are annotated by cell type using a combination of known gene markers and the available cell surface antibodies.

Upon running `nipalsMCIA` for $n = 10$ factors and plotting the global embedding of the samples for the first two factors, we observe that cells of the same type cluster together (Figure S2).

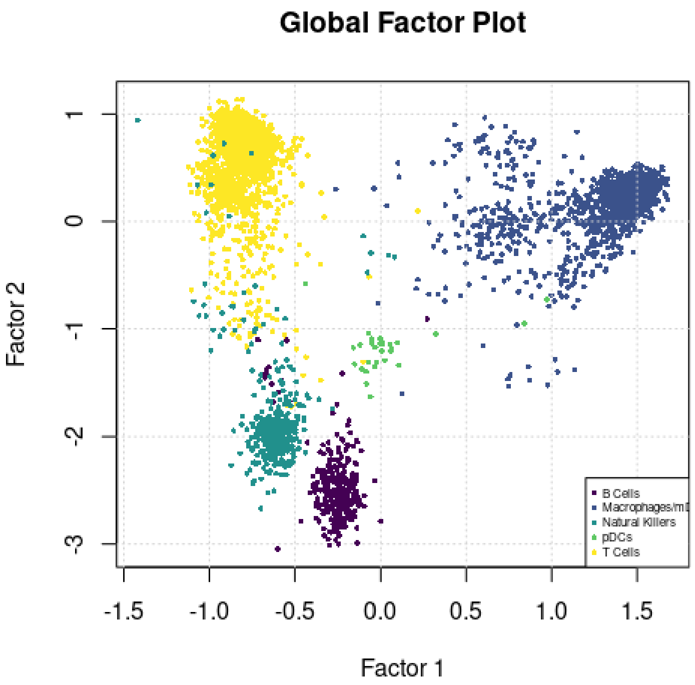


Figure S2: Global factor plot of the single cell data using the first two factors, colored by the annotated cell types (mDCs refers to myeloid dendritic cells, pDCs to plasmacytoid dendritic cells).

### References


[1] 10x Genomics. 5k peripheral blood mononuclear cells (pbmcs) from a healthy donor with cell surface proteins (v3 chemistry). May 2019.

[2] Abdifatah Abdi, Lynne Williams, Dominique Valentin, and Bennani-Dosse. Statis and distatis: Optimum multitable principal component analysis and three way metric multidimensional scaling. *Wiley Interdisciplinary Reviews: Computational Statistics*, 6:124–167., 03 2012.

[3] Daniel Chessel and M. Hanafi. Analyses de la co-inertie de k nuages de points. 1996.

[4] Mohamed Hanafi, Achim Kohler, and El-Mostafa Qannari. Connections between multiple co-inertia analysis and consensus principal component analysis. *Chemometrics and Intelligent Laboratory Systems*, 106(1):37–40, March 2011.

[5] Sahar Hassani, Mohamed Hanafi, El Mostafa Qannari, and Achim Kohler. Deflation strategies for multi-block principal component analysis revisited. *Chemometrics and Intelligent Laboratory Systems*, 120:154–168, 2013.

[6] Chen Meng, Bernhard Kuster, Aedín C. Culhane, and Amin Moghaddas Gholami. A multivariate approach to the integration of multi-omics datasets. *BMC Bioinformatics*, 15(1):162, May 2014.

[7] Chen Meng, Oana A. Zeleznik, Gerhard G. Thallinger, Bernhard Kuster, Amin M. Gholami, and Aedín C. Culhane. Dimension reduction techniques for the integrative analysis of multi-omics data. *Briefings in Bioinformatics*, 17(4):628–641, July 2016.

[8] Jonathon Shlens. A tutorial on principal component analysis. *International Journal of Remote Sensing*, 51, 04 2014.

[9] Age K Smilde, Johan A Westerhuis, and Sijmen De Jong. A framework for sequential multi-block component methods. *Journal of Chemometrics: A Journal of the Chemometrics Society*, 17(6):323–337, 2003.

[10] Grace X. Y. et al. Zheng. Massively parallel digital transcriptional profiling of single cells. *Nature Communications*, 8(1), January 2017.
